# Healed predation scar on a Cambrian apex predator

**DOI:** 10.64898/2026.08.01.742255

**Authors:** Kun-sheng Du, Yu Wang, Juan Gao, Stephen Pates, Wei Li

## Abstract

Predation is considered a key driver of the rapid diversification of animals during the Cambrian explosion. While the fossil record documents a plethora of evidence of successful and failed predation on biomineralized invertebrates at low trophic levels, no previous evidence of predation on larger, often soft-bodied, animals at higher trophic levels has been reported. This means that the modeled links between higher trophic levels in Cambrian food webs lack supporting fossil evidence, hindering understanding of the complexity of Cambrian trophic relationships. Here, we report a healed injury on the swimming flap of the radiodont apex predator *Amplectobelua symbrachiata*— one of the largest animals in the Cambrian oceans. The diagnostic W-shape with a healed margin supports interpretation of this wound as predatory in origin. Possible attackers include larger contemporaneous radiodonts – possibly members of the same species – or the giant lobopodian *Omnidens.* Interpreted as a healed predation scar, this W-shaped injury supports previous hypotheses of mutual predation on top predators in the Cambrian Stage 3, providing further evidence that trophic complexity increased during the latter stages of the Ediacaran-Cambrian Transition.

## INTRODUCTION

Predation is an important ecological driver of the Cambrian explosion (Wood and Zhuravlev, 2012), acting through predator preferences (e.g., Babcock and Robison, 1989; Pates et al., 2017) and predator-prey arms races (Bicknell et al., 2025) to shape evolution of both predators and prey. Healed predation scars on the biomineralized exoskeletons of small invertebrates, such as brachiopods, molluscs, tommotiids, and trilobites comprise the vast majority of evidence for predation in the Paleozoic (Bicknell and Paterson, 2018; Vinn, 2018), with much rarer direct evidence derived from coprolites and gut contents (e.g., Vannier and Chen, 2005; Zacaï et al., 2016; Bicknell and Paterson, 2018). Predation traces are observed chiefly on primary or secondary consumers including brachiopods, hyoliths, molluscs, and most trilobites (Dunne et al., 2008; Bicknell and Paterson, 2018; Vinn, 2018; Perrier et al., 2015; Park et al., 2024). Evidence for predation on animals at higher trophic levels (e.g., tertiary, quaternary) is limited to W-shaped predation traces on large paradoxidid trilobites (possible secondary or tertiary consumers; Bicknell et al., 2022; Mahata and Pates, 2026) and inferences from functional morphology and modeling suggesting that some predatory arthropods would prey on juveniles of other large arthropods (Dunne et al., 2008). This in turn means that the inferred modern ecological network structure in Cambrian marine ecosystems lacks a foundation in fossil data.

Here, we report a repaired injury to the flap of *Amplectobelua symbrachiata,* one of the largest swimming animals in Cambrian oceans at up to 90 cm in length (Wu et al., 2024a). The injured animal was likely around 23 cm in length. Interpreted as resulting from a predatory attack on an apex predator (quaternary consumers, e.g., Park et al., 2024), this specimen provides support from the fossil record of mutual predation among top predators in Cambrian ecosystems.

## MATERIALS AND METHODS

Specimens originate from the Yu’anshan Shale Member (or Yu’anshan Formation) of the Helinpu Formation (Chiungchussu Formation), *Eoredlichia*-*Wutingaspis* zone, Cambrian Series 2, Stage 3, Xincun section, Malong District, Qujing City, Yunnan Province (Figs. 1–4, 6A, and 6B). In previous studies, this stratum was typically assigned to the Chengjiang Biota (Hu, 2005). However, it is worth noting that *Eoredlichia*—an index fossil widespread across the Kunming and Yuxi localities of the Chengjiang Biota (Hu, 2005), the roughly coeval Fandian Biota in Sichuan (Du et al., 2020), and the contemporaneous Qingjiang Biota in Hubei (Fu et al., 2019)—has not yet been found in this interval from the Malong area. Material is accessioned at Research Center of Natural History and Culture, Qujing Normal University (XC-).

**Figure 1.**
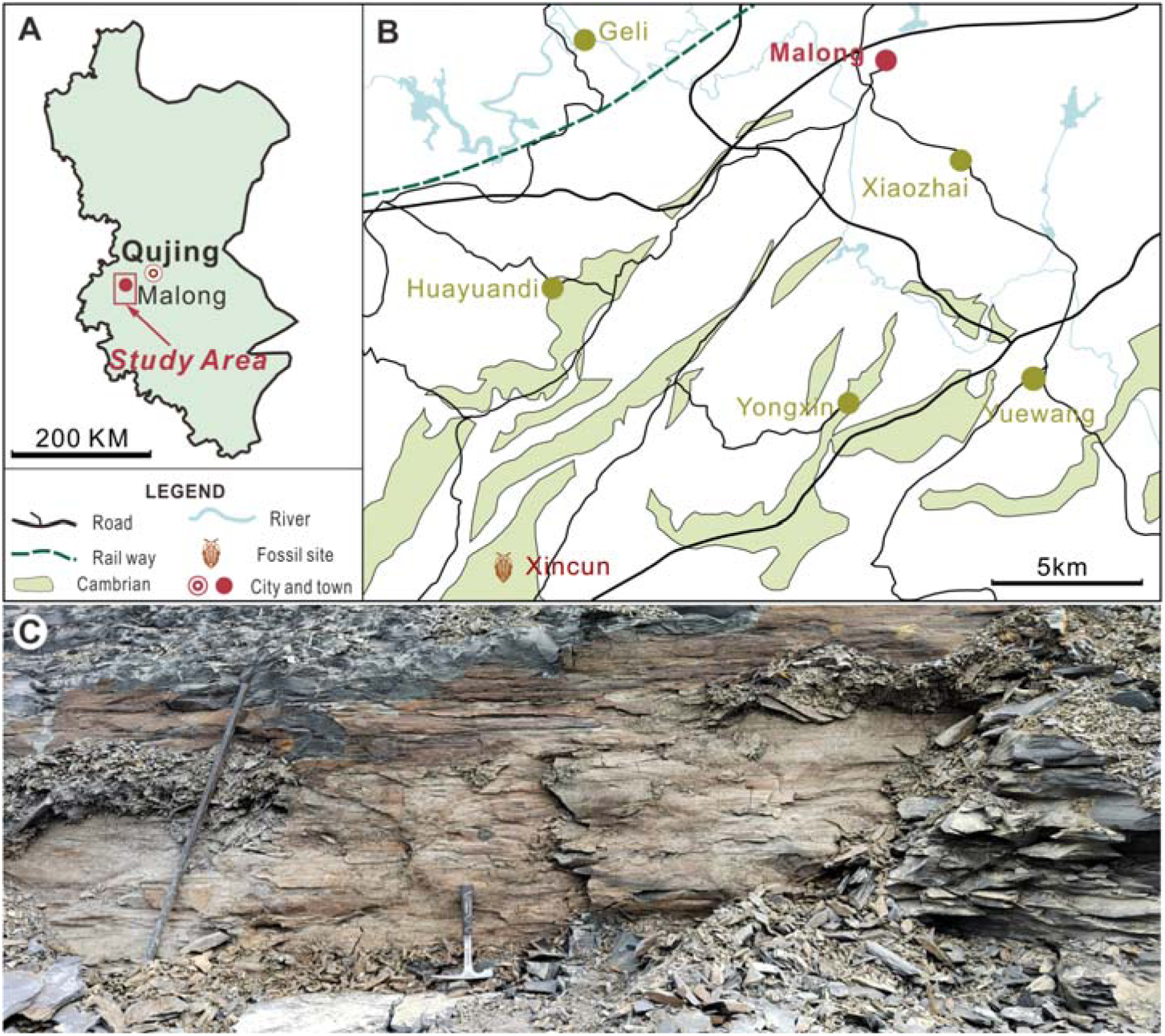
Locality maps of the Cambrian Stage 3 Xincun section and the strata yielding the flaps.

**Figure 2.**
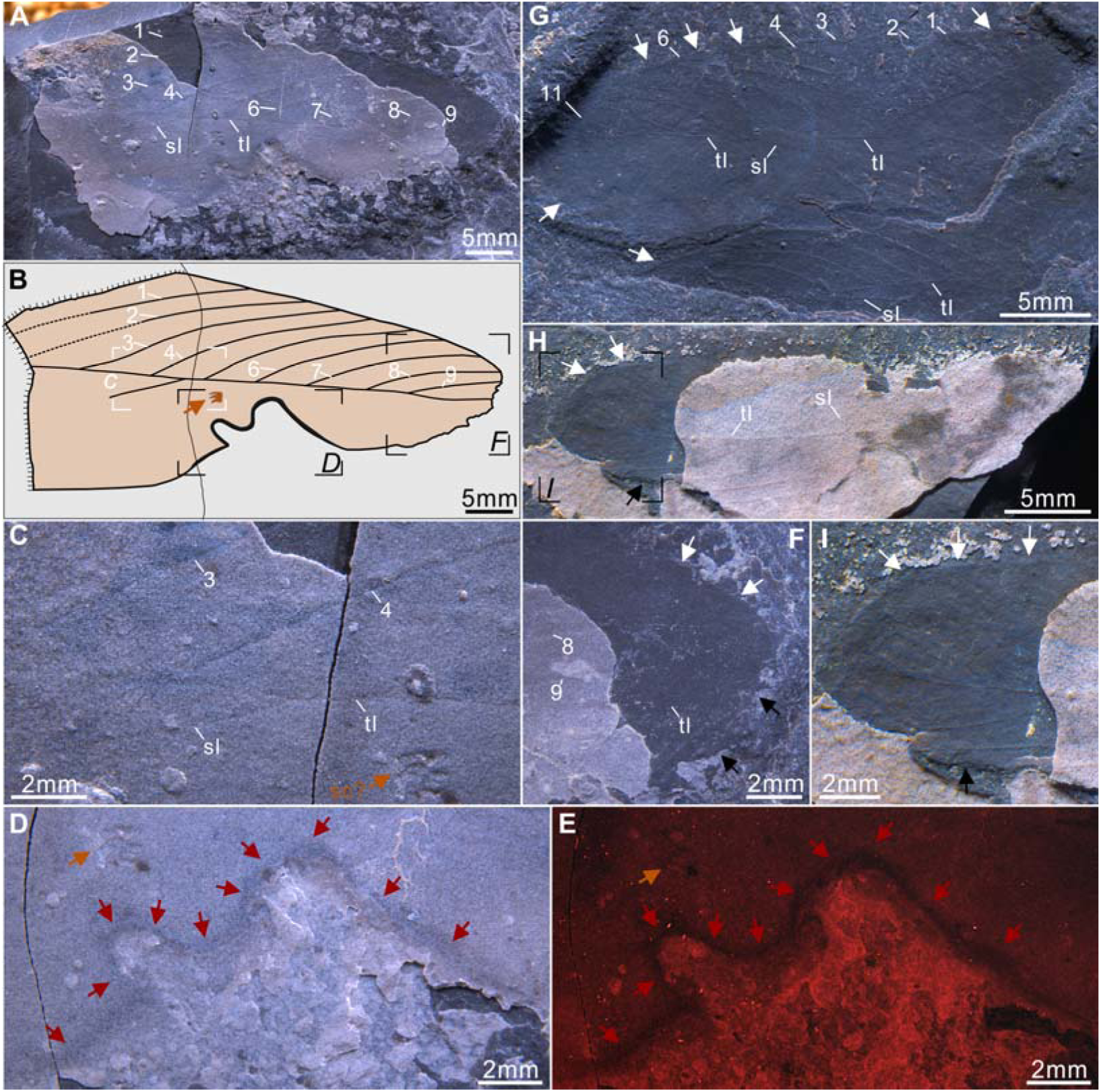
*Amplectobelua symbrachiata* flaps from Xincun section. (A to F), XC-018, a flap with transverse line (tl), nine oblique longitudinal lines (number 1–9), a short oblique posterior transverse line (sl), an irregular fold (sc?, orange arrow) and a W-shape healed injury with dark band of consistent thickness (red arrows). (G to I), XC-019, three flaps with preserved longitudinal line, up to 11 oblique transverse lines, and short oblique transverse line posterior to the longitudinal line. White arrows refer to unbroken margins of the flap, and black arrows to broken margins.

**Figure 3.**
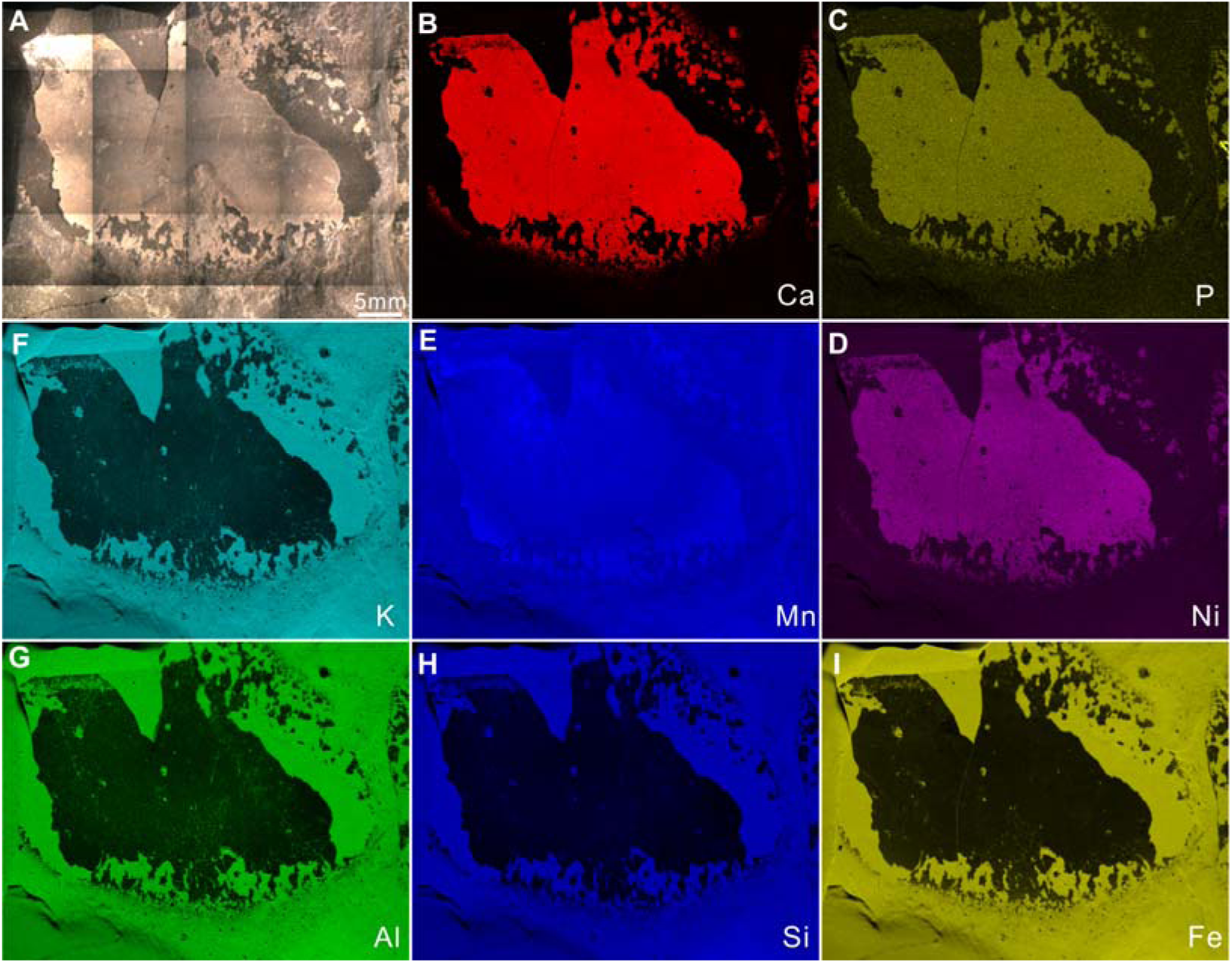
Micro-XRF elemental mapping results of the healed flap. (A), overview of the healed flap with thin coating. (B**–**E), showing the rich content of Ca, P, Ni, and Mn within the thin coating. (F and G), rich content of K, Al, Si, Fe within matrix.

**Figure 4.**
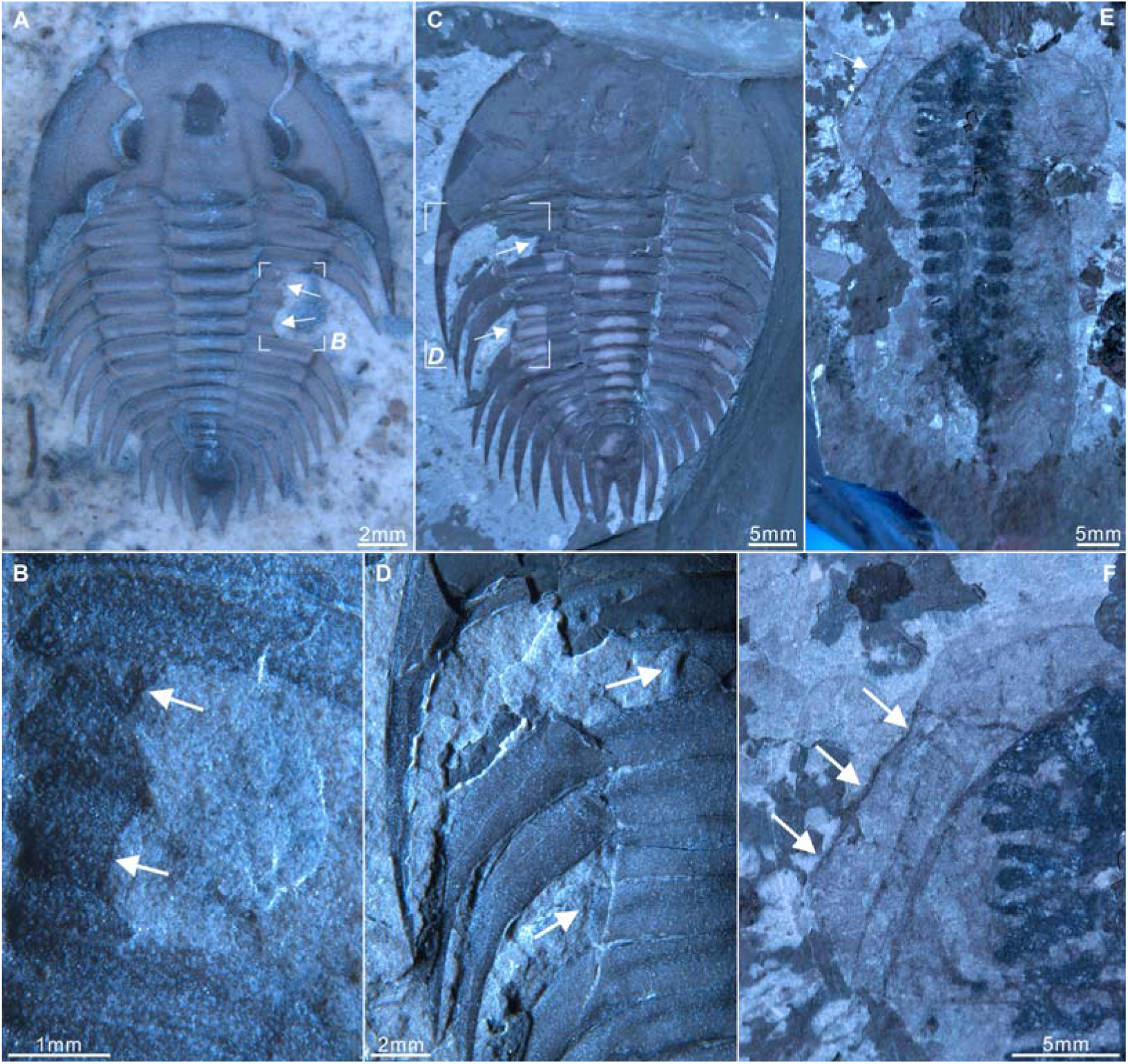
Healed injuries of euarthropods from Xincun. (A, B), XC-017a, *Malungia laevigata* bearing W-shaped injury (arrows). (C, D), XC-020, *M. laevigata* with W- and U-shaped injury. (E, F), XC-022, *Naraoia longicaudata* with gut diverticulae and healed injury (arrows). (A, C, E, F), wet and direct light; (B, D), dry and low light.

The specimens were photographed by a Leica DFC 550 digital camera mounted to a Stereoscope Leica M205 C, under different lighting conditions. The fluorescent images were captured using a Leica DFC7000 T monochrome camera attached to a Leica M205 FA fluorescence stereomicroscope. X-ray fluorescence elemental mapping was undertaken using an EDAX ORBIS PC XRF spectrometer. Explanatory drawings were prepared using CorelDraw X8, which was also used to prepare the figures.

## RESULTS

### Radiodont swimming flap with W-shaped embayment

The radiodont flap as the focus of this study measures 50.6 mm in length and 22.2 mm in width (Fig. 2A), giving a minimum body length for the whole animal of 23 cm (or larger if this flap was not the largest flap, based on Chen et al., 1994). The flap features nine oblique transverse lines on its anterior section and a single longitudinal line bisecting the flap into equal anterior and posterior portions (Figs 2A and 2B). The lines, preserved as negative relief in both the upper (lighter) and lower (darker) layer—exposed where the upper layer has cracked—taper distally (Figs 2A, 2B, and 2F). In contrast, transverse lines in the other specimens are preserved as positive relief (Figs 2G–2I). The posterior section of the flap contains only a single short, oblique line intersects the longitudinal line approximately one-third of the distance from the proximal end (Figs 2A–2C), this feature is also preserved on the other three flaps (Figs 2G and 2H). The W-shaped embayment is present near the middle of the posterior section (Figs 2A, 2B, 2D, and 2E), measuring 7.9 mm (shortest embayment), 15.8 mm (total width) and 9.6 mm (embayment on the right side). A distinct black strip of consistent thickness of 0.3 mm follows the edge of the W-shape, visible under both normal and fluorescence lighting (red arrows in Figs 2D and 2E). This strip is absent from broken or cracked margins elsewhere on the flap, and the darker layers are not exposed beneath the upper layer in this region, unlike in the anterior section. The margins (both broken and complete) of other radiodont flaps from this deposit lack any comparable black strip (Figs 2F–2I). An irregular fold occurs anterior and slightly more proximal to the W-shape (Figs 2B–2E, orange arrows).

### Taxonomic identification of the flap

This flap is attributable to *Amplectobelua symbrachiata* based on its size and arrangement of transverse lines. Transverse lines are diagnostic features of radiodonts (Potin and Daley, 2023) and are commonly observed in articulated radiodonts specimens and isolated flaps (Chen et al., 1994; Potin and Daley, 2023). Of contemporaneous radiodonts from the Chengjiang biota from which flaps have been described, *Innovatiocaris* has slender flaps with only seven transverse lines (Zeng et al., 2023). *Houcaris* (Hou et al., 1995), *Shucaris* (Wu et al., 2024b), *Lyrarapax* (Cong et al., 2016), *Ramskoeldia* (Cong et al., 2018), and *Anomalocaris* cf. *canadensis* (Wu et al., 2021a) have indeterminate transverse line counts, however, the angles between these transverse lines and flap anterior margins are different, and their transverse lines are closer to the front marginal rim than in our material. The most distinctive feature of our material is a short oblique transverse line in the posterior region bisected by the longitudinal line (Figs 2C, 2G, and 2H), which is only reported in the specimen ELRC 21001 of *A. symbrachiata* (see Figs. 3B and 4D in Chen et al., 1994). This identification is corroborated by the large size of the flap, as *A. symbrachiata* is one of the largest radiodonts from South China and from the Cambrian Stage 3 (Wu et al., 2024a), as well as by the overwhelming abundance of *A. symbrachiata* appendages at Xincun section relative to other radiodonts (over 800 isolated frontal appendages of this species have been collected to date, the next most abundant radiodont is *H. saron,* known from only three specimens).

## DISCUSSION

The morphology of the W-shaped feature, combined with comparisons to malformations in other fossil groups and consideration of the biology of *Amplectobelua symbrachiata*, supports our hypothesis that this represents a predation wound.

### This W-shaped embayment represents a repaired injury

The W-shaped embayment is best interpreted as a repaired injury due to its shape, evidence for healing, and comparison with broken margins along other parts of the fossil slab. It is unlikely to have been caused by decay or during the burial process.

The W-shape was neither caused by the decay process nor burial. X-ray fluorescence mapping (Fig. 3) shows that the coating of injury flap is rich in Ca, P, Ni, and Mn, whereas the matrix is dominated by Fe, Si, Al, and K, indicating a carbonate-containing coating and a silicate-rich matrix. Exquisitely preserved brachiopod fossils with ultrastructural details and diverse taphonomic pathways revealed apparent calcitic preservation as primarily iron oxides with subsequent coating (Huang and Rong, 2025), and this preservation style is similar to that of this flap and other specimens from Xincun (Figs 2, 4, 6A, and 6B). Damage from burial tends to exhibit irregular margins (black arrows in Figs 2F, 2H, and 2I), accompanied by sharp elemental compositional gradients between the matrix and the flap (Fig. 3).

Furthermore, centimeter and millimeter-scale structures such as this healed injury can be well preserved in thermal taphonomy experiments at 300°C (He et al., 2025), and the peak metamorphic temperatures of Cambrian Burgess Shale-type deposits from China are lower (Chengjiang ∼ 300 °C, Qingjiang ∼ 240 °C; Qiao et al., 2024). Thus, thermal alteration during burial is unlikely to have produced this W-shaped feature. Taphonomy experiments on marine arthropods demonstrate that decay can form irregular shapes (Krause et al., 2011). However, no similar W-shapes from Burgess Shale-type deposits have been attributed to the decay process, with these shapes normally interpreted as injuries (Babcock and Robison, 1989; Bicknell and Paterson, 2018; Bicknell et al., 2022).

The W-shaped embayment to the flap (red arrows in Figs 2D and 2E) exhibits a dark thickened strip indicative of healing, analogous to cicatrized scars in trilobites (Figs 4A–4D, e.g., Owen, 1985; Mahata and Pates, 2026) and other arthropods (Figs 4E and 4F), and consistent with the melanisation (Fig. 5) present during appendage healing in extant arthropods (Bilandžija et al., 2017) and implies that the damage occurred and was healed while the animal was alive. This flap may have been shed during ecdysis, as the likely non-lethal predation affected less than 30% of its posterior section, constituting only minor trauma relative to the whole body. As only this one flap is preserved, it is not known if multiple flaps were damaged, or if the attack was limited to just this one flap.

**Figure 5.**
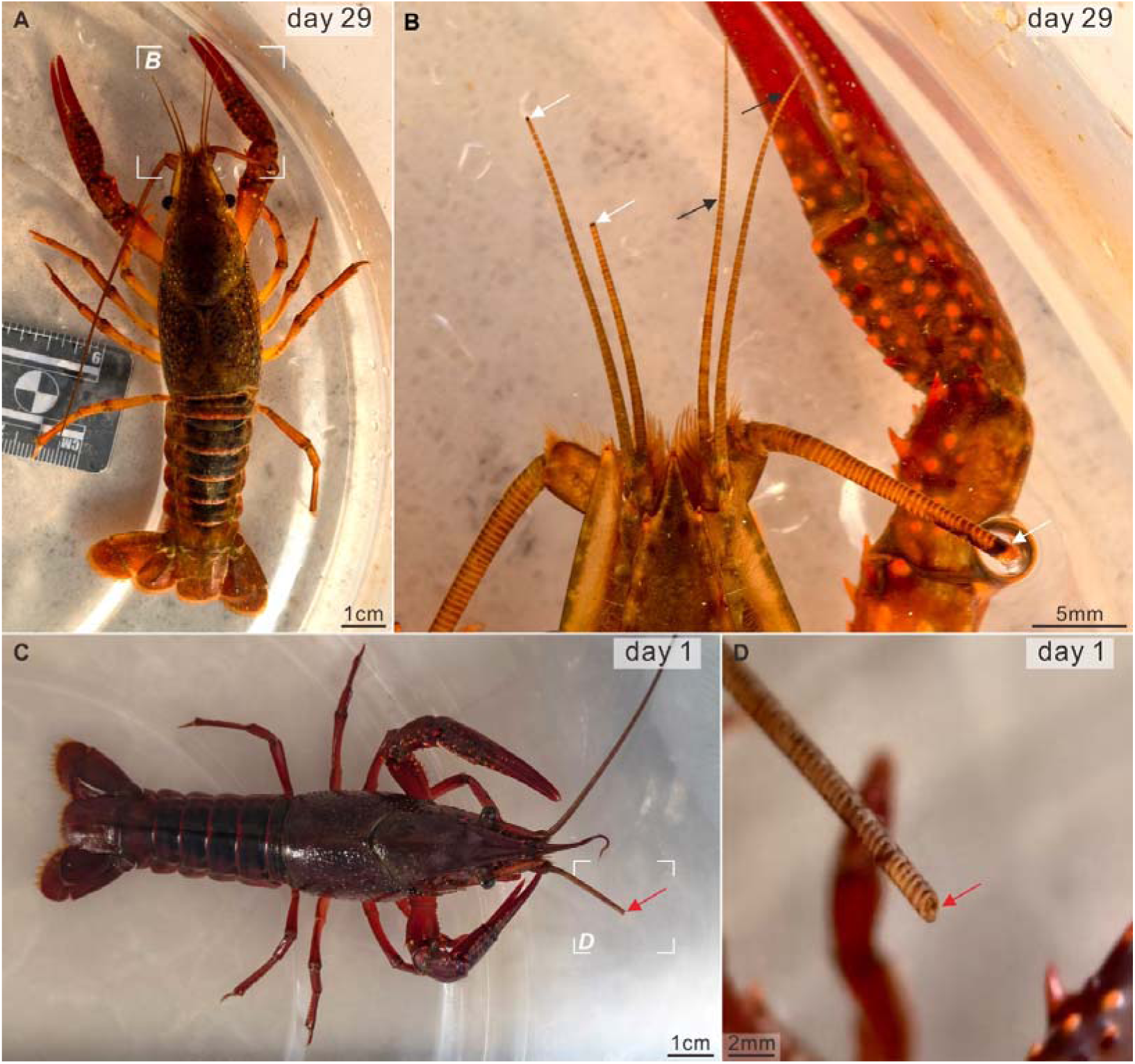
**Injuries on appendages of modern arthropod *Procambarus* sp**. (A, B), melanization in the repaired appendages after 29 days (white arrows), and corresponding normal appendages (black arrows). (C, D), new injury without obvious melanization (red arrows).

### Repaired injury is interpreted as healed predation wound

From a sample size of one, we cannot apply statistical approaches as used in studies of shelly prey to support injuries as predatory in origin (e.g., Leighton 2011; Harper 2016). In studies of Cambrian trilobites, preference for a particular location (e.g., right side; Babcock and Robison 1989; posterior; Pates et al., 2017, Bicknell et al., 2022), and/or size (Bicknell et al., 2022) has been used to support a predatory interpretation for injuries in studies with large sample sizes. However, in the extensive literature on trilobite malformations and abnormalities, W-shaped outlines showing evidence of repair are interpreted as predatory in origin (Figs 4A–4D, e.g., Bicknell and Paterson, 2018). The W-shape and evidence of repair, and comparison with the extensive record of repaired injuries on trilobite exoskeletons (Figs 4A–4D), provide support for interpreting this W-shaped embayment as a repaired injury from predation, rather than alternative explanations that lack support.

In studies of abnormalities in trilobites, generally three main causes are considered: injuries from predation or molting; parasite induced pathologies; and developmental teratologies (Owen 1985; Babcock 1993). Arthropods are commonly injured during molting, however these injuries are expected to occur in areas where there is a narrow feature that tears during the exit from the exoskeleton – such as a genal spine in a trilobite (e.g. Owen 1985; Losso and Bicknell 2026) – rather than the broad U-shaped swimming flap of a radiodont. Parasite induced pathologies and developmental teratologies tend to produce local atrophy and unusual growths and lack evidence of repair (e.g., Owen 1985; Bicknell et al., 2018). Evidence of repair and the W-shape support the identification of this embayment as a repaired injury, while its shape and location on a feature not expected to be difficult to extract during the molting process support predation rather than molting as the primary cause. The relative sizes of wound-maker and wounded *A*. *symbrachiata* provide additional support for the predation hypothesis, since if the wound was caused by a predatory attack, the predator would be expected to be larger than its prey. Estimates of the prey and predator size—derived from the sizes of the flaps and the spacing of endites or oral cone parts, based on the size of the wound (see Supplemental Material)—suggest that the wound-maker was at least 50% larger, and possibly well over 100% larger than the injured radiodont.

Rarely, injuries to fossil arthropods have been attributed to sexual selection and intraspecific combat, for example in the trilobite *Walliserops* (Gishlick and Fortey, 2023). In *Walliserops*, a prominent trident is compared to the horns of beetles, and interpreted as a sexually selected character. No such supporting evidence for radiodonts exists, and no features have been suggested to be sexually selected for any member of this group. Therefore, we do not have a foundation to infer that the W-shaped injury was caused during sexual selection or competition for mates. Similarly, there are limited reports of dominance hierarchies in marine arthropods (e.g., Rappaort and Lord, 2021; Lord et al., 2021). Support for a dominance hierarchy origin for the W-shaped injury in *Amplectobelua* could come from larger size in a ‘dominance tool’ (the chelipeds of grass shrimp, or the frontal appendages in *Amplectobelua*) relative to body size in some individuals compared to others.

Alternatively, perhaps the enlarged proximal endite in some individuals could have been larger than others, given its importance in making a ‘pincer’. Only one specimen of *A*. *symbrachiata* with appendages and body preserved together is known (Chen et al., 1994, fig. 3), while the appendages of *A. symbrachiata* grew isometrically (Wu et al., 2024a), meaning that we cannot support hierarchical competition hypothesis from morphological evidence. Alternative or complementary evidence could come from identifying a limited resource in Xincun, for which hierarchical competition would provide a benefit (Chase et al., 2002). From what we know of the function of *Amplectobelua* appendages, they could grasp in a ‘pincer’ (Daley and Budd 2010; De Vivo et al., 2021). Radiodonts were nektonic swimmers, and even if we consider *A. symbrachiata* also limited to non-biomineralized prey like *Anomalocaris canadensis* (e.g., Daley and Bergstrom 2012; Bicknell et al., 2023), there were a diverse array of prey animals in Xincun, including many species that were abundant. For example, three species of small lobopodian, and more than 3 species of non-biomineralizing euarthropods including *Naraoia longicaudata* and *Combinivalvula chengjiangensis* each known from over 1000 specimens so far. There is therefore no supporting evidence for hierarchical competition being favorable in Xincun, or morphological support for hierarchical competition in *A*. *symbrachiata*.

Thus, the best interpretation for the W-shaped healed wound is as a repaired injury from predatory attack, as this interpretation has support from comparison to numerous other W-shaped injuries in trilobites that have been assigned to predation.

If this interpretation is accepted, the W-shaped injury represents the first reported repaired predation trace on a radiodont, or any large Cambrian non-biomineralized predator. Evidence of repaired injury to soft-bodied animals in the Cambrian is extremely rare due to the low preservation potential of these fossils (known from far fewer sites than those hosting biomineralized fossils), and is compounded by the difficulties in demonstrating repair (rather than other types of damage) in soft-bodied material. While biomineralized exoskeletons generally exhibit clear outline, and often three-dimensional preservation with even cicatrization of injuries known (Owen, 1985), these features are much less likely to be clear in flattened carbonaceous films common to Burgess Shale-type fossils. Thus, this specimen with evidence for a repaired margin, a diagnostic W-shape to the damage, and clear distinction between broken and repaired margins, is extremely rare.

### Who hunted hunters in Cambrian oceans?

Identifying the predator responsible for injuries recorded on fossil specimens is often very challenging (e.g., Harper 2016). The size, shape, and location of the repaired injury on the flap allow us to suggest some possibilities. Potential predators of *Amplectobelua* must have appendages and/or mouthparts suited for active predation (Figs 6 and 7) and sufficient body size to cause an injury of this size. A W-shaped injury on the flap could be produced through the interaction of the enlarged distal dorsal spines, terminal spines, the hypertrophied endite on the first claw podomere, and endites of alternating length along the appendage (Fig. 6C). Of Chengjiang radiodonts with this appendage morphology, only those of *A. symbrachiata* (>135 mm), *Ramskoeldia* (∼150 mm) or *Laminacaris* (∼280 mm) approach the requisite size (Cong et al., 2018; Guo et al., 2019; Wu et al., 2024a). The estimated wound-maker likely bore a frontal appendage ∼70 mm in length or mouthpart ∼88 mm in diameter (Supplemental Material). Other radiodonts from Chengjiang with raptorial appendages are too small to have created such a large injury. Those of *Innovatiocaris* reached ∼32 mm (Zeng et al., 2023)*, Lenisicaris* and

**Figure 6.**
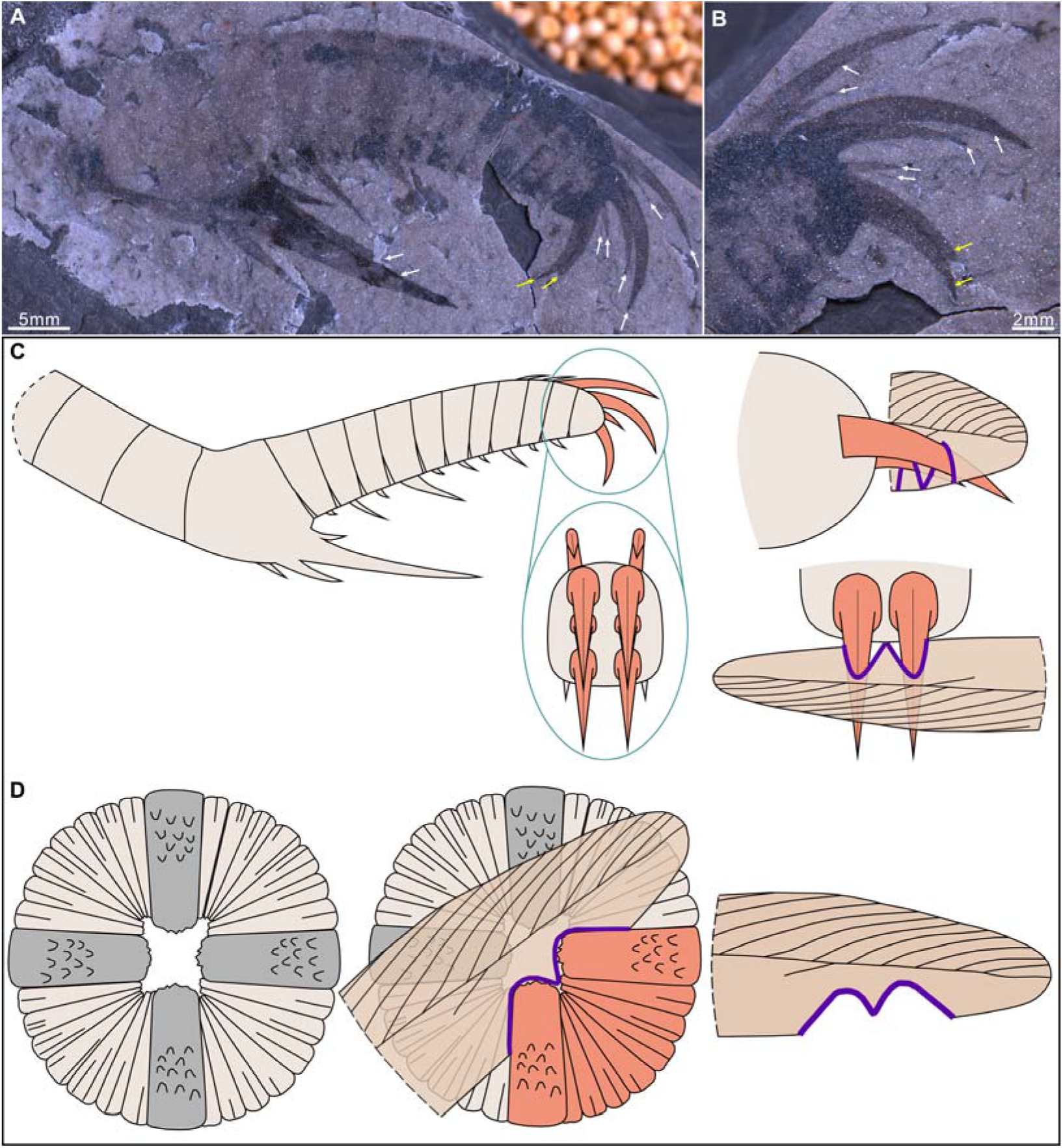
W-shaped injuries caused by radiodonts. (A, B), XC-029, *Amplectobelua symbrachiata* frontal appendage with paired spines (arrows), note the paired terminal spines (yellow arrows). (C), mechanism of W-shaped injury by a frontal appendage. (D), mouthpart producing the W-shaped injury.

**Figure 7.**
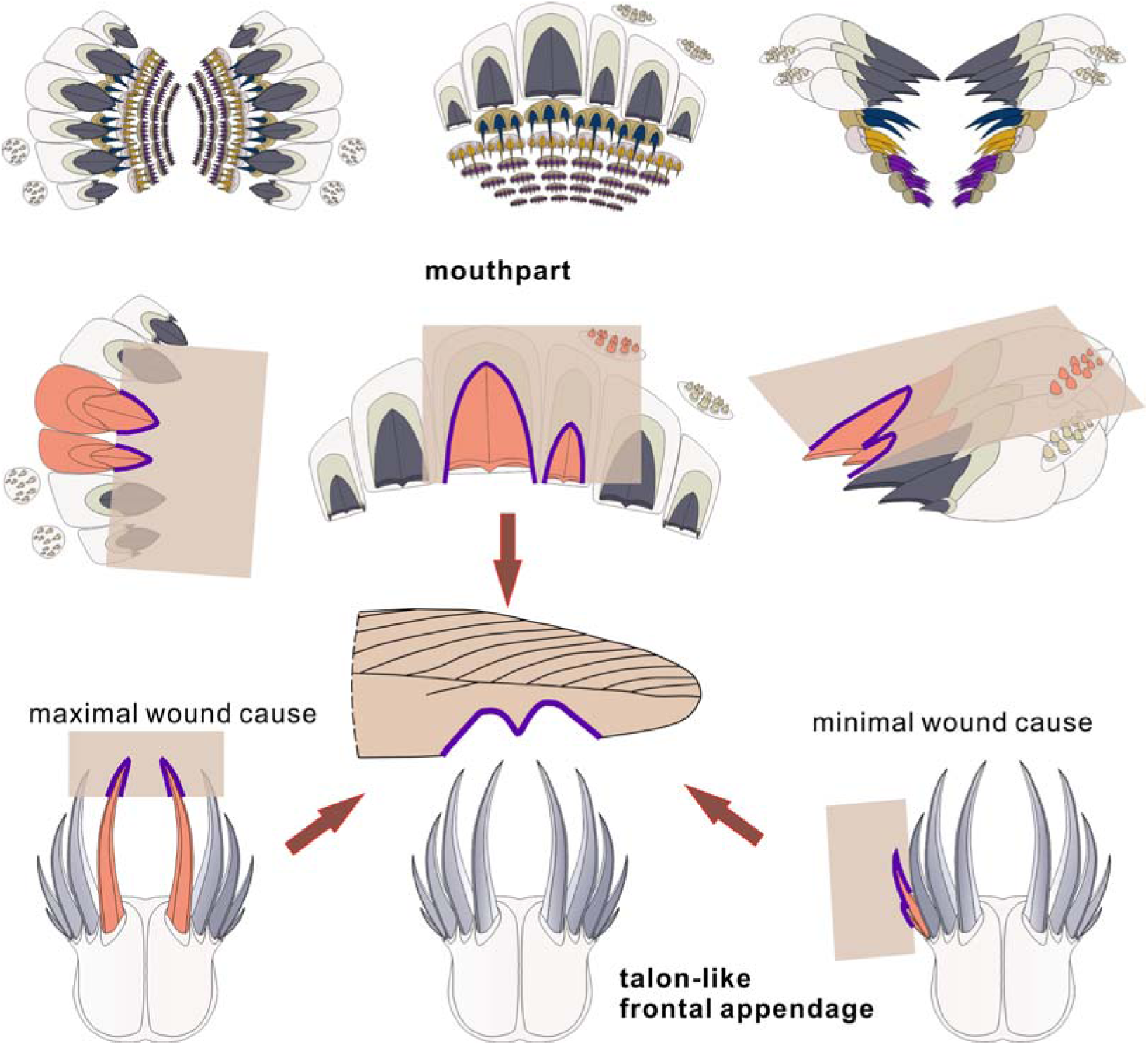
Schematic diagram of hypothesized W-shaped injuries caused by *Omnidens*.

*Lyrarapax* ∼25 mm (Cong et al., 2016; Wu et al., 2021a), *Shucaris* ∼40 mm (Wu et al.,

2024b), while hurdiid radiodonts (Wu et al., 2022) and those with fine spines such as *Houcaris* (Wu et al., 2021b) lack of suitable morphology for producing a W-shaped injury. Similarly, only large radiodonts such as *A. symbrachiata*, *Ramskoeldia* and *Laminacaris* would have had large enough oral cones to cause comparable damage. An oral cone is not known in *Amplectobelua,* and so any inferences on oral cone size and morphology are based on the closely related *Ramskoeldia.* The biomechanics of radiodont oral cones are currently not well understood, but recent hypotheses have suggested that they were adapted for suction feeding rather than mastication (e.g., Daley and Bergström, 2012; Potin et al., 2026). If an oral cone caused the damage to this flap, the prey animal would have been brought to the cone by the appendages.

Gnathobase-like structures, known in both *A. symbrachiata* and *Ramskoeldia* (Cong et al., 2017, 2018) are unlikely to have caused this injury due to their smaller size (Supplemental Material). Apart from radiodonts, the contemporaneous giant predator *Omnidens* could also have inflicted similar injuries with its talon-like frontal appendages and/or mouth apparatus (Fig. 7). This animal has large mouthparts with a length of ∼123 mm, composed of multiple toothed plates (Fig. 4F in Zhang et al., 2014). Any two principal plates consist of one large and one small component, consistent with the left-small and right-large configuration of the W-shaped injury (Fig. 7). The irregular fold (orange arrow in Figs 2B–2E) could be compared to the toothed plates of *Omnidens*, and may be a remnant of the bite that produced this W-shaped injury (Fig. 7).

### Implications for Cambrian food webs

If the W-shaped injury is interpreted as a predation wound, regardless of the exact predator, this would provide support from the fossil record for previous inferences that apex predators were attacked within Cambrian food webs (Dunne et al., 2008). These modeled Cambrian ecosystems were inferred to have had a comparable complexity to modern marine ecosystems (Dunne et al., 2008). Given that *A. symbrachiata* itself represents a possible wound-maker, Cambrian cannibalism, commonly observed in modern marine species like crabs (Hines et al., 2026), is also of potential ecological significance. Cannibalism and short high trophic-level feeding loops are common in modern food webs across a range of ecosystems (Hines et al., 2026), and here we suggest the presence of cannibalism at the highest trophic level in the Cambrian (Fig. 8), in an ecosystem from over half a billion years ago, complementing reports of potential cannibalism in, and predation on, large Cambrian trilobites (Bicknell et al., 2022; Mahata and Pates, 2026) possibly secondary or tertiary consumers. Whether due to cannibalism or another large contemporaneous predator, if interpreted as a predation injury, this specimen provides support for previously hypothesized increase in food web complexity from the Ediacaran to the Cambrian. Relatively simple and linear Terreneuvian and Ediacaran webs (Mitchell and Pates, 2025), became more comparable to food webs in the modern ocean from the Cambrian Stage 3 with higher trophic levels feeding on one another (Dunne et al., 2008), albeit with panarthropods and chaetognaths rather than vertebrates at the highest trophic levels (Dunne et al., 2008; Perrier et al., 2015; Park et al., 2024).

**Figure 8.**
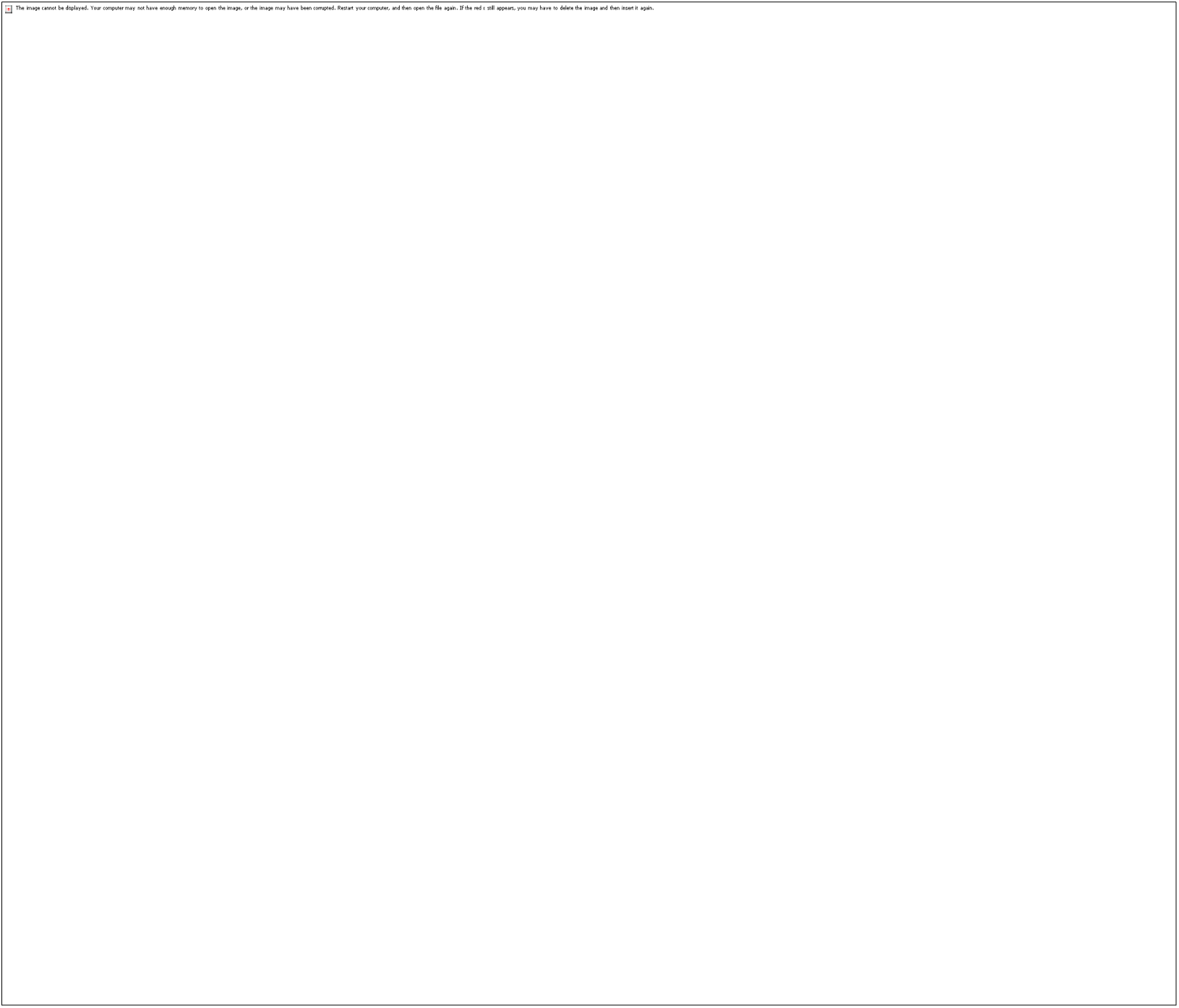
Cambrian trophic levels. Predation on *Amplectobelua* indicating the presence of possible cannibalism at the highest level. Red arrows indicate possible cannibalism. Revised from Perrier et al. (2015), and Park et al. (2024).

## CONCLUSION

The first evidence of a Cambrian apex predator with a healed wound is a swimming flap with a W-shaped embayment. Morphological comparisons to similar wounds in trilobites support interpretation of this wound as predatory in origin, providing support from the fossil record that the largest apex predators were hunted within Cambrian ecosystems.

## ACKNOWLEDGMENTS

This work was supported by the National Natural Science Foundation of China (grant numbers 42662002, 42202003, and 42262004 to K.S.D.), NERC IRF NE/X017745/2 (S.P.), and the Caiyun Postdoctoral Program Innovation Project of Yunnan Province (W.L.). We are very grateful to Professor Q. Ou (China University of Geosciences, Beijing) for help in field work and for his identification of the strata. Constructive criticisms by L. Lustri and J. Shaw and two anonymous reviewers greatly improved the manuscript.

## 1. Calculation

### *Body size of the injured* Amplectobelua symbrachiata

According to the relatively complete specimen of *A. symbrachiata* (Fig. 3A in Chen et al., 1994), the lateral flaps exhibit a decreasing size trend towards the posterior of the body. The first five pairs of flaps are relatively large, and their distal portions would have been more susceptible to grasping by predators. In contrast, the remaining flaps show a clear trend of posterior contraction, rendering them less vulnerable to capture. Therefore, it is reasonable to estimate the size of this individual based on the dimensions of the first five pairs of flaps.

On the third right flap (Fig. 3A in Chen et al., 1994), the transverse lines are relatively well preserved. Measurements indicate that the length from the intersection of the third transverse line with a longitudinal line near the tip to the tip itself is approximately 1 cm. This represents a ratio of about 1:12 relative to the total body length (excluding frontal appendages and furcae). The corresponding distance in this study (Fig. 2A) measures approximately 2.1 cm.

Measurement of the length from the flap tip to the trunk midline shows that the ratio between the third right flap and the fifth left flap is approximately 4:3. This ratio indicates that, for the fifth flap, the distance from the aforementioned intersection point to its tip constitutes about 6.25% of the total body length. Conversely, the ratio between the third right flap and the first right flap is approximately 1:1.1, suggesting that first flap accounts for about 9.17% of the trunk length.

Therefore, applying the corresponding length from Fig. 2A of this study to this range (6.25%–9.17%) suggests that the length of the prey *A. symbrachiata* was approximately 22.9–33.6 cm. Considering measurement uncertainty, the estimated length is roughly 23–35 cm.

### Estimated predator size from frontal appendages

Given an appendage-to-body ratio of ∼1:6 (appendage measured from the base of the first claw podomere to the distal tip), the prey *A*. *symbrachiata* had 3.8–5.8 cm long appendages. As predators typically exceed prey in size (Donadio and Buskirk, 2006), and only three contemporary Cambrian taxa are known with larger appendages (Cong et al., 2018; Guo et al., 2019; Wu et al., 2024a): *A. symbrachiata* (>135 mm), *Ramskoeldia* (∼150 mm) and *Laminacaris* (110–280 mm). The latter two are known only from isolated appendages or lack sufficiently complete soft-bodied preservation for reliable total body length estimation; moreover, they belong to different genera.

Therefore, the similar wounds are attributed to predation rather than intraspecific interaction. In *A. symbrachiata*, the articulation between claw podomeres and arthrodial membranes restricts the appendage to grasping prey and delivering it to the mouth (De Vivo et al., 2021). Consequently, the only appendage elements capable of grasping the flap are the paired dorsal spines and the terminal spines (Figs 6A–6C). The maximum wound size indicates the minimum predator size. Measuring the largest dorsal spine gap (on claw podomere 11) from Chen et al. (1994), yields ∼0.5 cm, ∼8.3 % of appendage length. With a minimum wound separation of 0.58 cm on the flap (Fig. S1), the calculated predator body length is 0.58 ÷ 8.3% × 6 ≈ 42 cm.

Therefore, the attacking *A*. *symbrachiata* was ≥ 42 cm long, with ≥ 7 cm long frontal appendages.

### Estimation of predator size based on oral cone

Although biting force generated by the oral cone remains unknown (Nedin, 1999; Daley and Bergström, 2012), the elements that produce the largest W-shaped wound are the adjacent large plates and the small plates situated between them (Fig. 6D).

This attribution provides a basis for estimating the minimum size of the predator.

*Laminacaris* known only from appendages and its phylogenic position is uncertain (Guo et al., 2019), thus, its oral cone size cannot be inferred from wound dimensions. Although no oral cone has been described for *A. symbrachiata*, other amplectobeluid species—namely, *Ramskoeldia*, *Lyrarapax* and *Guanshancaris*—are known to possess a tetradial oral cone (Liu et al., 2018; Jiao et al., 2021; Wu et al., 2024b). Thus, these results provide a provisional basis for estimating the oral cone of

*A. symbrachiata*.

In a tetradial cone, the opening made by plates varies in outline (Daley and Bergström, 2012), yet all plates share a common center. Modelling the cone as a set of concentric circles minimizes error. The vertical distance from the lowest point of the wound (blue dot in Fig. S1) to the flap margin far exceeds its distance to the two highest points (green and orange dots in Fig. S1), indicating that the small plates also capture the flap. As struggling prey and variable contact angles between the plates and the flap can alter the spacing of the two high points; therefore, the distance between a high point and its adjacent low point was used as the reference. The largest possible wound—and thus the minimum predator size—occurs when the tip of a large plate and the tip of the smallest plate act together at 45° angle. Using concentric circles and isosceles triangle geometry, the oral cone size can be calculated. Since the oral cones of the contemporaneous amplectobeluids *Ramskoeldia* and *Lyrarapax* are incomplete preserved (Liu et al., 2018; Wu et al., 2024b), but the lengths of their large and smallest plates can be measured, the ratio (La-b/D) of the difference in plate lengths (La-b) to oral-cone diameter (D) serves as a proxy. For *Lyrarapax*, La-b/D ≈ 4.8% (Liu et al., 2018); for *Ramskoeldia*, La-b/D ≈ 5.2% (Wu et al., 2024b). *A. symbrachiata* is expected to fall within this range. With a measured wound La-b ≈ 4.6 mm (Fig. S1), the corresponding oral cone diameter would be 4.6 mm ÷ 4.8% to 4.6 mm ÷ 5.2% ≈ 8.8–9.5 cm for *A. symbrachiata*, or ∼8.08 for *Ramskoeldia*.

Given that the cephalic region constitutes ∼10% of the trunk length in *A. symbrachiata* (Fig. 3A in Chen et al., 1994), and the oral cone is approximated by the cephalic region (a conservative assumption, as the cone is slightly smaller), the estimated oral cone diameter (8.8–9.5 cm) yields a total predator body length of (8.8– 9.5 cm) × 10 ≈ 88–95 cm.

Gnathobase-like structures have a maximum width of 18 mm and bear four distal spines. The two adjacent distal spines reach a maximum length of ∼8.4 mm (Cong et al., 2018; Wu et al., 2024b), indicating that they are too small to produce the W-shaped injury with a width of 15.8mm. If three or more distal spines caused the injury, the wound would not be W-shaped. Therefore, gnathobase-like structures are excluded from the present size estimation.

## 2. Supplemental Figure

**Figure S1.**
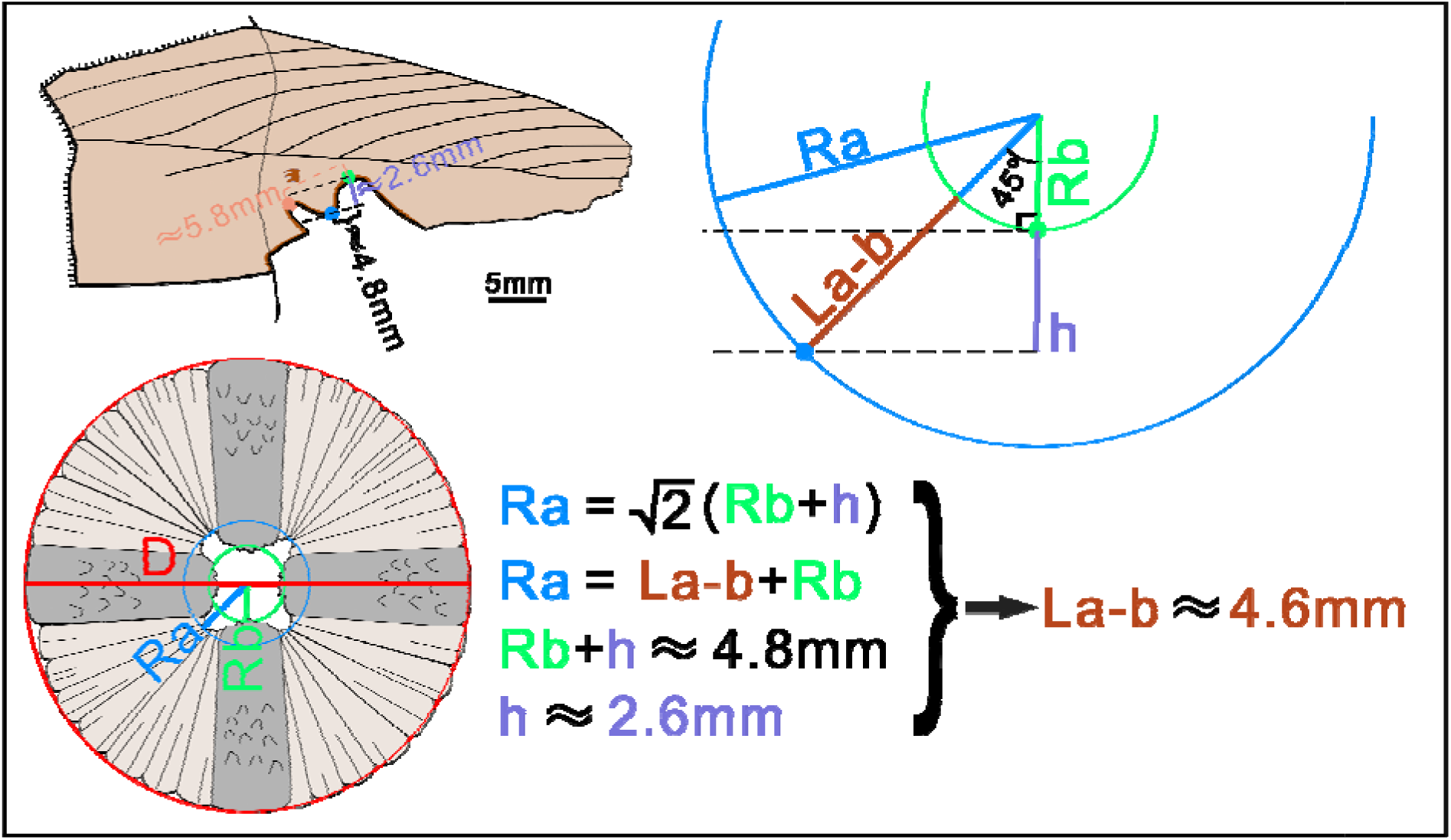
Measured wound data and calculative steps based on oral cone.

